# Control over patch encounters changes foraging behaviour

**DOI:** 10.1101/2021.01.19.426950

**Authors:** Sam Hall-McMaster, Peter Dayan, Nicolas W. Schuck

## Abstract

Foraging is a common decision problem in natural environments. When new exploitable sites are always available, a simple optimal strategy is to leave a current site when its return falls below a single average reward rate. Here, we examined foraging in a more structured environment, with a limited number of sites that replenished at different rates and had to be revisited. When participants could choose sites, they visited fast-replenishing sites more often, left sites at higher levels of reward, and achieved a higher net reward rate. Decisions to exploit-or-leave a site were best explained with a computational model that included both the average reward rate for the environment and reward information about the unattended sites. This suggests that unattended sites influence leave decisions, in foraging environments where sites can be revisited.

**Highlights:**

- Being able to select sites during foraging increased visits to high value sites
- This visitation pattern was efficient, producing higher average reward rates
- Decisions to leave a site were influenced by information about alternative sites

**Graphical Abstract:** 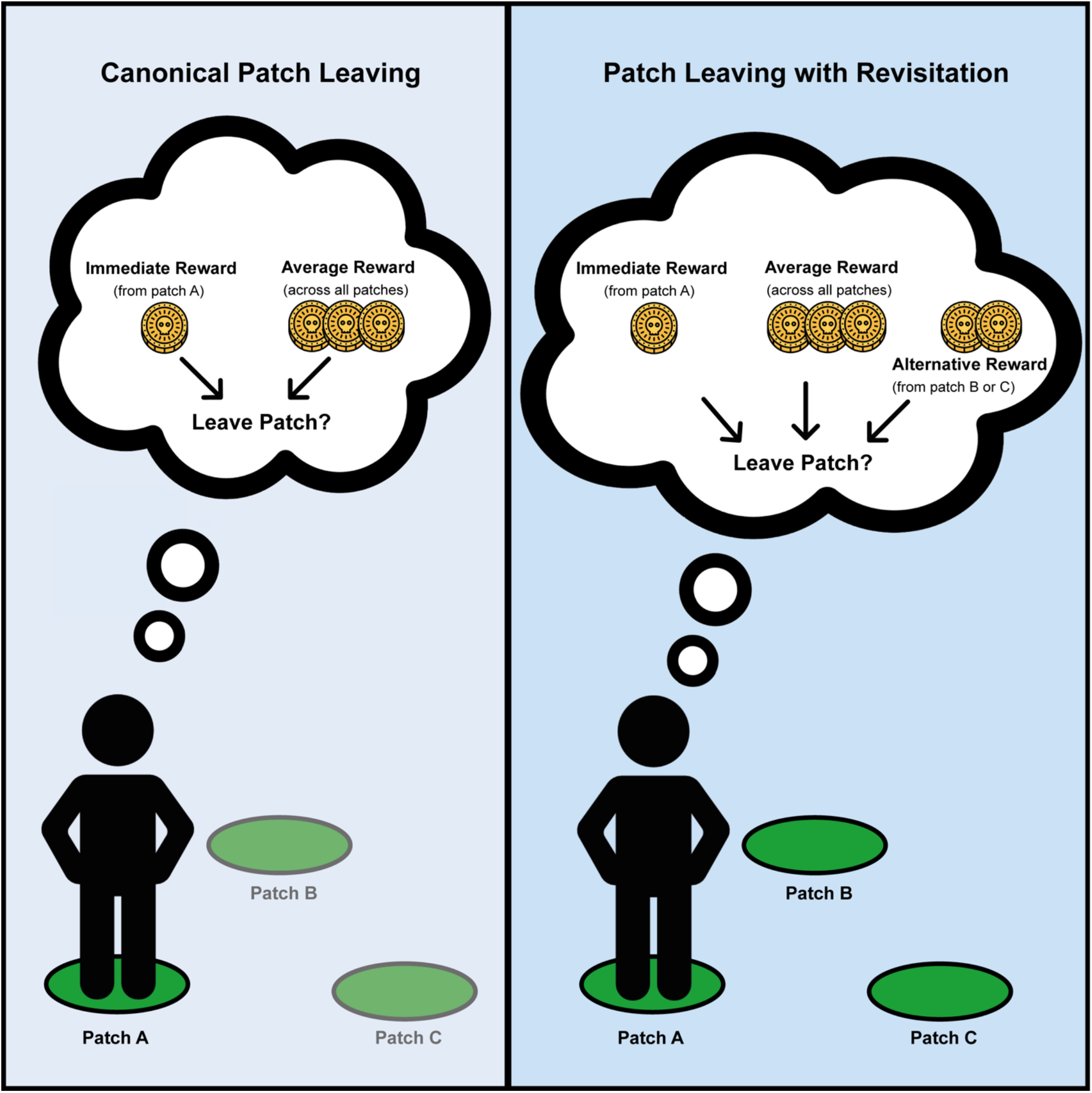

## Introduction

Decision making requires anticipating how our choices will influence our fortunes in the future. Much work in neuroscience has therefore focused on how decision making can be understood as an optimisation problem based on a Markov Decision Process (MDP) formalism, in which future environmental states and rewards are under partial control of the decision maker (Sutton & Barto, 1998). This approach can account for behaviour in complex environments, where planning is needed to form an optimal policy (Huys et al., 2012; Huys et al., 2015; Kurth-Nelson et al., 2016), and can explain neural activity found in dopaminergic areas during value prediction (Schultz et al., 1997).

A common decision problem for animals in natural environments is foraging, in which it is often assumed that animals have limited control over future environmental states. During a particular kind of foraging called patch-leaving, animals must decide between exploiting a current patch or leaving to search for a better alternative. Canonical Optimal Foraging Theory (OFT) considers a simplified version of this problem, in which patches within an environment are encountered at fixed rates (Charnov, 1976). Under these conditions, the average reward rate in an environment is the sole variable needed to anticipate future reward outcomes for deciding to leave. This results in a simple optimal policy called the Marginal Value Theorem (MVT, Charnov, 1976), wherein animals should leave their current patch when its reward falls below the average reward rate for the environment. Empirical studies have shown that leave decisions in mice, rats, monkeys and humans are consistent with this decision rule (Constantino & Daw, 2015, Hayden et al., 2011; Kane et al., 2017; Le Heron et al., 2020; Lottem et al. 2018). Evidence has also suggested that the unique structure of foraging decisions might be solved using neural substrates that are at least partially distinct from those involved in other reward-based decisions (Kolling et al., 2012; Rushworth et al., 2011), with the anterior cingulate cortex serving a critical role in regulating leave decisions (Fouragnan et al., 2019; Hayden et al., 2011; Wittmann et al., 2016, see Kolling et al., 2016; Mobbs et al., 2018; Rushworth et al., 2011 for reviews).

While canonical foraging decisions might not require the full algorithmic complexity of MDP solutions and could be solved using distinct brain areas from other forms of decision making (Kolling et al., 2012; Rushworth et al., 2011), more general foraging choices might not be as simple as MVT assumes. For example, patches that animals encounter are not just determined by their frequency in an environment, but through decisions animals make about which patches are worth exploiting (Kamil, 1978; Merkle et al., 2014; Passingham, 1985; Sayers & Menzel, 2012; Sweis et al., 2018). In addition, resources in natural environments can replenish following exploitation (Berger-Tal & Avgar, 2012; Garcia & Eubanks, 2019), making it valuable to revisit specific sites at later time points (Erwin, 1985, Lihoreau et al., 2010; Merkle et al., 2014; Seidel & Boyce, 2015). This introduces a new dilemma about when return is appropriate that is not considered under MVT because it assumes little to no revisitation (Charnov, 1976; Possingham & Houston, 1990). Behavioural ecology research has long been aware of these issues, as evidenced by studies examining the contributions of memory to foraging decisions in environments where patches can be revisited (Berger-Tal & Avgar, 2012; Merkle et al., 2014), and have often approached the problem of how animals store information in memory for future behaviour using reinforcement learning (RL) and Bayesian updating (Berger-Tal & Avgar, 2012; Marshall et al., 2013). These additional foraging considerations raise important questions about what drives leave decisions when encounters with patches are under the decision maker’s control and when patches promise different rewards due to their growth dynamics. Are decisions to leave an option still based on global reward information, such as the average reward rate across options in the environment, or are these choices made using local reward information about specific alternatives?

We recently proposed that foraging-inspired tasks used in neuroscience could be made more ecological by allowing animals to choose which patch is encountered, after deciding to leave the current patch (Hall-McMaster & Luyckx, 2019). In the present experiment, we therefore studied foraging decisions in an environment where foraging choices could be directed towards specific patches, and in which patch resources replenished over time. Compared to situations where options are encountered at random, we predicted that decision makers would use their knowledge about patch differences to visit faster replenishing patches more often, increasing the average reward per action. We also predicted that decisions to leave a patch would be made earlier, when a patch had higher reward levels and had been exploited fewer times. Based on MVT, this change in leaving threshold would be expected due to a higher global reward rate. However, an alternative mechanism would be that decision makers leave based on option-specific reward estimates when their next option can be selected.

In the present study, we tested the predictions above and sought to address how global reward information (about all patches) and local reward information (about specific patches) are used to decide when to leave a patch. Human participants completed a patch-leaving task similar to studies from Optimal Foraging Theory (Stephens & Krebs, 1986), in which a patch’s reward decreased over time through exploitation. In contrast to previous studies (Constantino & Daw, 2015, Hayden et al., 2011; Kane et al., 2017; Le Heron et al., 2020), participants switched between the same three patches throughout a block, and patches replenished their rewards at distinct rates when not being exploited. Participants therefore needed to decide when to leave their current patch in order to revisit and exploit one of the two alternatives. Each block involved 200 actions. Actions included selecting a patch to visit, exploiting it for reward and leaving it. Across blocks, we manipulated whether participants had a free choice over which patch to exploit after deciding to leave the current patch or whether the choice was forced, being randomly determined for them.

When participants were able to control what patches were encountered, we found that fast replenishing patches were visited more often than slow replenishing patches. This increased the average reward rate and elevated leaving thresholds, with higher reward outcomes prior to leave decisions. Due to higher reward both when arriving at patches and when leaving them, no difference was observed in the number of exploit decisions before leaving, between free and forced choice conditions. Participants’ exploit-or-leave decisions were best explained with a computational model that included both the global reward rate and local information about the unattended patches.

## Results

To investigate the effect of decision control on foraging behaviour, 60 participants completed an online foraging task. Participants controlled a pirate ship, sailing to three equidistant islands to dig for buried treasure. Participants first selected an island to sail to and then made a series of exploit-or-leave decisions. When deciding to exploit, participants dug for treasure and received between 0 and 100 gold coins based on the island’s current reward level. The number of coins depleted super-exponentially across successive exploit actions on the island. While the current island was being exploited, new treasure was buried at each alternative island at an island-specific rate (slow, medium or fast). When deciding to leave the current island, participants sailed to one of the two alternative islands. This structure repeated until participants had performed a total of 200 actions with the same three islands, and the block ended. Choosing islands to visit, exploiting islands and leaving all counted as individual actions. Sailing did not consume any actions in the block.

The central manipulation in this task was whether participants had a free or forced choice over the island they would sail to next, after deciding to leave the current island. In free choice blocks, the seas were calm and participants could select either alternative island, following a leave decision. In forced choice blocks, the seas were stormy and islands could not always be reached. Participants were therefore forced to sail to just one of the alternative islands, determined at random, following each leave decision. The alternative island that participants would be forced to sail to was only revealed after each leave decision was made.

Participants completed two free choice blocks and two forced choice blocks in a random order, and were informed about the upcoming condition prior to each block. Islands were located at one of three vertex positions that formed an equilateral triangle on screen. Slow, medium and fast replenishing islands were assigned at random to different vertex positions for each block. Participants were told that blocks would contain a fast, slow and medium replenishing island before starting the experiment, but did not have direct experience or practice with the islands before starting. In the results that follow, we refer to each island as a patch, consistent with terms used in Optimal Foraging Theory (Stephens & Krebs, 1986; Charnov, 1976).

### Participants directed visits in proportion to expected patch rewards

We first examined how often participants decided to visit each patch under free and forced choice conditions. In line with our predictions, we found a significantly higher proportion of visits to the fast replenishing patch in free compared with forced choice blocks (mean free choice=0.366, SD=0.027; mean forced choice=0.328, SD=0.037; corrected *p*=1e-4), and a significantly lower proportion of visits to the slow replenishing patch in free choice blocks (mean free choice=0.289, SD=0.035; mean forced choice=0.341, SD=0.035; corrected *p*=1e-4, see Figure 2A). The proportion of visits to the medium replenishing patch was also significantly higher in free choice blocks (mean free choice=0.345, SD=0.025; mean forced=0.331, SD=0.037; corrected *p*=0.018). All proportions, both here and hereafter, were compared using non-parametric permutation testing (see STAR Methods), in which corrected *p*-values were calculated based on the rank of the true *t*-statistic within a null distribution generated from 10,000 random data permutations. In cases where the true *t*-statistic was higher than all values in the null distribution, the corrected *p*-value becomes one over the number of permutations (i.e. 1e-4).

Exploratory analyses revealed that participants selected the alternative patch with the higher expected reward (i.e. current patch value) significantly more under free choice conditions, as compared to forced choice conditions, wherein participants were forced to select a patch determined at random after each leave decision. This effect was observed when leaving the slow patch (mean free=0.711, SD=0.195; mean forced=0.511, SD=0.121; corrected *p*=1e-4), the medium patch (mean free=0.615, SD=0.178; mean forced=0.480, SD=0.113; corrected *p*=1e-4) and the fast patch (mean free=0.641, SD=0.166; mean forced=0.493, SD=0.135; corrected *p*=1e-4, see Figure 2C). When considering choices based on patch replenishment rates, rather than current patch values, participants selected the alternative patch with the higher replenishment rate significantly more when leaving the medium replenishing patch under free choice conditions (mean free choice=0.615, SD=0.171; mean forced choice=0.455, SD=0.134; corrected *p*=1e-4, see Figure 2D). To a lesser extent, this was also the case when leaving the fast replenishing patch (mean free choice=0.570, SD=0.189; mean forced choice=0.510, SD=0.112; corrected *p*=0.036). Significant differences were not detected in how often the higher replenishment rate patch was selected when leaving the slow replenishing patch (mean free choice=0.539, SD=0.196; mean forced choice=0.510, SD=0.125; corrected *p*=0.355).

### Reward rates were higher when patch encounters could be directed

The ability to direct foraging towards patches with faster replenishing dynamics should result in higher rewards overall and, therefore, increase experienced reward rates. In line with this prediction, we observed a significant main effect of choice condition on the average reward per exploit action (*F*(1,59)=40.526, *p*=3.170e-8, see Figure 3). There was also a significant main effect of patch type (*F*(2,118)=275.161, *p*=3.670e-45) and a significant interaction between choice condition and patch type (*F*(2,118)=12.234, *p*=1.485e-5). The main effect of choice condition was due to higher reward rates in the free compared to forced choice blocks (mean free choice=68.268, SD=9.221; mean forced choice=64.119, SD=8.538). The main effect of patch type was due to increasing reward rates from the slow (mean=62.202, SD=9.257) through to the medium (mean=67.025, SD=8.391) and fast replenishing patch (mean=69.382, SD=8.220). The interaction between choice condition and patch type reflected decreasing differences in the reward rate between choice conditions as the replenishment rate of a patch increased from slow to medium (mean difference slow=5.601, SD=6.540; mean difference medium=3.864, SD=5.565; *t*(59)=-3.645, corrected *p*=0.002) and slow to fast (mean difference fast=3.094, SD=4.497; *t*(59)=-4.231, corrected *p*=2.473e-4) but not from medium to fast (*t*(59)=-1.664, corrected *p*=0.304). The increase in reward rate under free compared with forced choice conditions was significantly different from zero for all three patch types (slow *t*(59)=-6.635, corrected *p*=3.357e-8; medium *t*(59)=-5.379, corrected *p*=-4.048e-6; fast *t*(59)=-5.239, corrected *p*=6.810e-6). Including non-exploit actions in the reward rate calculation lowered the average reward across choice conditions (mean free choice=47.510, SD=2.662; mean forced choice=45.409, SD=2.668) and patches (mean slow=41.684, SD=2.582; mean medium=47.362, SD=2.555; mean fast=50.332, SD=2.683) but did not change the pattern of results above.

### Reward threshold for leave decisions was higher when patch encounters could be directed

Optimal Foraging Theory (Charnov, 1976) predicts that as experienced reward rates increase, patches will be abandoned at higher reward values. The previous result, showing higher reward rates in free compared to forced choice blocks, therefore suggests there might be differences in the average reward value before leaving. To test this idea, we examined the last reward outcome before leaving each patch under free and forced conditions. We observed a significant main effect of choice condition (*F*(1,59)=18.723, p=5.923e-5), reflecting higher reward outcomes prior to leave decisions in free (mean=48.753, SD=17.903) compared with forced choice blocks (mean=44.418, SD=16.001, see Figure 4A). There was also a significant main effect of patch type (*F*(2,118)=15.376, *p*=1.164e-6), reflecting lower reward outcomes before leaving the fast replenishing patch (mean=45.503, SD=16.547), compared to the medium replenishing patch (mean=46.728, SD=16.970; *t*(59)=-3.481, corrected *p*=0.003) and compared to the slow replenishing patch (mean=47.526, SD=16.329, *t*(59)=-5.009, corrected *p*=1.583e-5). A significant difference was not detected between the reward before leaving the medium and slow replenishing patches (*t*(59)=-2.322, corrected *p*=0.071). No significant interaction was detected between choice condition and patch type on the reward outcome prior to leaving (*F*(2,118)=0.021, *p*=0.980).

### Directed patch choices resulted in higher rewards on arrival and a comparable number of exploit actions before leaving

If participants selected fast replenishing patches more often, higher reward rates might not just reflect patches being abandoned at higher reward values, but the average patch reward being higher at arrival. When investigating this possibility, we found a significant main effect of choice condition on reward at arrival to a patch (*F*(1,59)=316.847, *p*=2.127e-25), indicating that the reward upon arrival at a patch differed in free and forced choice blocks (see Figure 4D). Moreover, we observed a significant main effect of patch type on reward at arrival (*F*(2,118)=675.677, *p*=2.402e-65) and a significant interaction between choice condition and patch type (*F*(2,118)=33.187, *p*=3.671e-12). Exploratory analyses revealed that the interaction resulted from a smaller differences in the arrival reward between free and forced choice conditions when arriving at the fast patch (mean difference=1.27, SD=2.12) compared to the medium patch (mean difference=2.697, SD=2.967; *t*(59)=2.807, corrected *p*=0.020), and when arriving at the medium patch compared to the slow patch (mean difference=6.552, SD=4.545; *t*(59)=5.364, corrected *p*=4.281e-6).

A related set of analyses, examining arrival rewards as a function of the patch being left, showed a significant main effect of choice condition on the arrival reward (*F*(1,59)=153.055, *p*=4.928e-18), reflecting higher arrival rewards in the free choice condition (mean free=80.470, SD=2.320; mean forced=76.036, SD=3.485, see Figure 4C). There was also a significant main effect of the patch being left on the reward gained when arriving at a new patch (*F*(2,118)=60.296, *p*=9.105e-19). Exploratory analyses showed that this main effect was due to higher arrival rewards after leaving the slow compared to the medium patch (mean slow=81.966, SD=2.950; mean medium=77.643, SD=3.972; *t*(59)=6.605, corrected *p*=3.770e-8), the slow compared to the fast patch (mean fast=75.150, SD=4.406; *t*(59)=10.975, corrected *p*=2.106e-15) and the medium patch compared to the fast patch (*t*(59)=4.102, corrected *p*=3.822e-4). No significant interaction was detected between the patch being left and choice condition (*F*(2,118)=1.185, *p*=0.310).

The combined effects of higher rewards at arrival and before leaving in the free compared to the forced choice condition resulted in a similar number of exploit actions before patch-leaving across conditions (mean free=5.251, SD=1.772; mean forced=5.452, SD=1.643; *F*(1,59)=3.337, *p*=0.073, see Figure 4B). There was a significant main effect of patch type (*F*(2,118)=336.641, *p*=1.736e-49) and a significant interaction between patch type and choice condition on the number of exploit actions before leaving (*F*(2,118)=9.312, *p*=1.758e-4). Exploratory analyses revealed that the interaction resulted from significantly fewer exploit actions prior to leaving the fast replenishing patch in the free choice condition (mean free choice=5.672, SD free choice=1.753; mean forced choice=6.015, SD forced choice=1.554; *t*(59)=3.400, corrected *p*=0.004), but no condition differences in leaving times for the slow replenishing patch (mean free choice=4.773, SD free choice=1.798; mean forced choice=4.759, SD forced choice=1.787; *t*(59)=-0.103, corrected *p*>0.99) or the medium replenishing patch (mean free choice=5.309, SD free choice=1.850; mean forced choice=5.581, SD forced choice=1.708; *t*(59)=2.207, corrected *p*=0.094).

Together with our analyses of arrival reward rates, these results suggest that when the reward at arrival was more closely matched across choice conditions (i.e. in the fast replenishing patch), patches were abandoned after fewer actions under free choice conditions. When the reward at arrival was not well matched across choice conditions (i.e. in the slow and medium replenishing patches), the higher rewards at arrival and prior to leaving seen in free choice blocks cancelled each other out, resulting in a similar number of actions before leaving to the forced choice condition.

### Local reward information influenced exploit-or-leave decisions

The results above indicate that being able to control patch encounters altered patch-leaving decisions. Fast and moderate replenishing patches were chosen more frequently, increasing reward rates in the free choice condition. Crucially, although higher reward rates would result in leaving thresholds according to MVT (Charnov, 1976), this could also be due to a change in computational mechanisms underlying leave decisions. For instance, participants could have not only used global reward rate information to guide their patch-leaving choices; they could have also used local information about patch-specific values or reward rates.

We tested this idea using logistic regression. First, we constructed a model that predicted exploit-or-leave decisions using the last reward outcome and the global reward rate across all patches, similar to the core variables used in MVT. We will call this the *global* model. Next, we added local reward variables to the global model, to test whether local information improved the model’s predictions in each condition. The global model was constructed as follows:

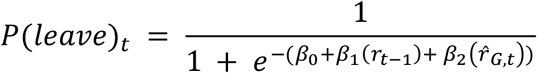

The core elements of this expression include *r*_*t*−1_, the last reward received from the current patch at time *t-1*, and 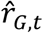, the average reward rate across all patches at time *t. β*_0_ is an intercept that estimates participant’s bias to leave, regardless of the last reward and the reward rate. *β*_1_ and *β*_2_ are estimated coefficients that weight the predictor variables *r*_*t*−1_ and 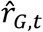. The average reward rate regressor, 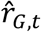, was estimated at time *t* using a delta learning rule, which updated the previous estimate using the last reward outcome, *r*_*t*−1_, and a learning rate, *α*:

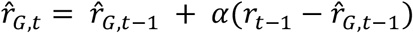

Given participants received no immediate reward for deciding to leave or selecting the next patch, 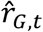 was updated with rewards of 0 following each of these actions. Since reward and no reward might influence participants’ estimate of the reward rate to different extents, we allowed the model to have separate learning rates for rewarded and unrewarded updates (*α* and *α*_*l*_). At the beginning of each block, the reward rate was initialised using a free parameter, *s*. While this approach models the probability of leaving at time *t*, leave events, and the corresponding predictor values were only observed if participants had chosen to exploit until time *t*-1. Hence, the approach used here can also be considered as model of the hazard function of leaving.

The *global+maxValue* model took the global model and included the maximum current value among the alternative patches, *r*_*v,t*_, as an additional predictor. The predictor value of *r*_*v,t*_ for each trial reflected the true state of the task (i.e. the reward participants would receive if they were to switch to the best possible patch). The static maximum reward rate model (*global+maxRR*_*S*_) was similar, except that it included the maximum reward rate among the alternative patches 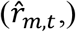 rather than the maximum of their current values. One important difference was that the additional regressor in the *global+maxValue* model increased over time, mimicking patch replenishment. In contrast, the additional regressor in the *global+maxRR*_*S*_ model remained static while exploiting the current patch. Trial-wise values for 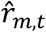 were obtained by estimating the reward rate separately for each patch *p*:

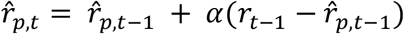

The learning rule above was used to update the reward rate estimate for the current patch *p*, but left the reward rate estimates of the other patches, *q*, unchanged:

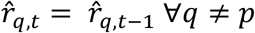

The local reward rates were updated with the same learning rates for rewards and no-rewards, and were initialised in the same manner as the global reward rate. The maximum of the two estimated reward rates of the two alternative (i.e. unattended) patches was then taken as the local predictor value on each trial. A central feature of the *global+maxRR*_*S*_ model was that it did not update the reward rates for unattended patches, which meant the maximum alternative reward rate was static while the current patch was being exploited. Given participants knew patches were replenishing over time, however, we created a more dynamic reward rate model (*global+maxRR*_*D*_), which updated reward rate estimates for unattended patches. For each action in the current patch, each alternative patch increased its reward rate towards its arrival reward (*r*_*A*_, see Figure 4D), from the last time that patch was visited:

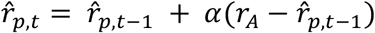

The highest reward rate among the alternatives was again used as the predictor value on each trial. All regression models used 3 free parameters to estimate the reward rate/s, *s, α* and *α*_*l*_. This meant that the same learning rates were used to estimate the global reward rate and the patch specific reward rates in the *global+MaxRR*_*S*_ and *global+MaxRR*_*D*_ models. All models used an additional 3-4 free parameters in the logistic regression equation, one for the constant term, *β*_0_, and one for each predictor, *β*_*n*_.

The results of the model comparison are shown in Figure 5. When comparing model performance in the free choice blocks, we did not detect consistent evidence that the *global+maxValue* model outperformed the global model on average (mean AIC/BIC difference=1.20/-2.38, negative difference values indicate the global model provided a better average fit to the choice data). When comparing the models in individual participants, we found that adding local value information to the global model resulted in better exploit-or-leave predictions for 25/60 participants based on AIC scores (10/60 participants based on BIC scores).

In contrast, there was consistent evidence that the *global+maxRR*_*S*_ model improved predictions in free choice blocks over the global model (mean AIC/BIC difference=5.38/1.80, positive difference values indicate the local model fit the data better on average). Individual subject analyses showed that 40 out 60 participants had lower AIC scores for the *global+maxRR*_*S*_ model (26/60 based on BIC).

Finally, when evaluating the *global+maxRR*_*D*_ model, which assumed reward rates were updated for unattended patches, we found mixed evidence that predictions were improved on average (mean AIC/BIC difference=3.34/-0.25). Individual subject analyses provided more compelling evidence for this model than average difference scores, showing that its predictions were better than the global model for 37/60 participants based on AIC scores (18/60 based on BIC).

When evaluating AIC scores across all four models, 23 participants were best fit by the *global+maxRR*_*S*_ model, 18 by the *global+maxRR*_*D*_ model, 10 by the global model and 9 by the *global+maxValue* model. We therefore concluded that the winning model for exploit-or-leave choices in the free choice condition was the *global+maxRR*_*S*_ model.

Despite the behavioural differences between choice conditions, we found a similar pattern of modelling performance for the forced choice blocks. The *global+maxRR*_*S*_ model outperformed the global model (mean AIC/BIC difference=7.56/3.96), providing the best fitting model overall for 22 participants based on the AIC and 15 participants based on the BIC. The *global+maxValue* model also showed a consistent performance improvement (mean AIC/BIC difference=4.02/0.43). However, its AIC/BIC difference scores were lower than the *global+maxRR*_*S*_ model and it explained fewer participants overall (12 based on the AIC and 8 based on the BIC). The *global+maxRR*_*D*_ model showed mixed evidence of improved performance relative to the global model on average (mean AIC/BIC difference=3.33/-0.26), providing the best fit for 23 participants based on the AIC and 11 based on the BIC. Considering both AIC and BIC scores, we concluded that the most consistent evidence pointed to the *global+maxRR*_*S*_ as the winning model in the forced choice condition.

The standardised coefficients for the winning *global+maxRR*_*S*_ model (Table 1) included a negative intercept for both choice conditions, indicating a bias to stay in patches regardless of the reward predictors. The highest coefficient seen in both conditions was the previous reward outcome, suggesting the instantaneous reward information had the largest relative influence on decisions to exploit-or-leave. The standardised coefficient for the local reward rate (*maxRR*_*S*_) was numerically higher than the global reward rate coefficient in both conditions. However, within-condition differences between these coefficients were not significant when scrutinised with permutation testing (Forced choice: corrected *p*=0.416; Free choice: corrected *p*=0.901; based on 10,000 random data permutations). In a similar manner, the difference between local and global coefficients was numerically higher in the free choice condition, compared to the forced choice condition. However, the difference was not significant (corrected *p*=0.496, based on 10,000 random data permutations). Parameters used to estimate the reward rates (*s, α* and *α*_*l*_) were numerically higher in the free choice condition.

**Table 1.** Median parameter estimates for the winning exploit-or-leave model under free and forced choice conditions. Bracketed values show the interquartile range. *β* values are standardised coefficients. *β*_0_ is the intercept, *β*_1_ is the previous reward coefficient, *β*_2_ is the global reward rate coefficient, and *β*_3_ is the maximum alternative reward rate coefficient. *s, α* and *α*_*l*_ are the starting value and learning rate parameters used to estimate the reward rates.

| Model | Condition | $\beta_0$ | $\beta_1$ | $\beta_2$ | $\beta_3$ | $s$ | $\alpha$ | $\alpha_l$ |
| --- | --- | --- | --- | --- | --- | --- | --- | --- |
| *Global+maxRR<sub>s</sub>* | Forced | -4.01<br>(3.03) | -4.51<br>(3.37) | 0.64<br>(1.85) | 0.73<br>(1.54) | 51.5<br>(26.45) | 0.25<br>(0.54) | 0.01<br>(0.18) |
|  | Free | -3.66<br>(2.5) | -4.25<br>(2.8) | 0.26<br>(2.68) | 0.79<br>(1.47) | 56.46<br>(27.15) | 0.33<br>(0.47) | 0.09<br>(0.19) |

## Discussion

Foraging has often been studied as a case of decision making in which animals have little control over patches encountered in the future (Barack et al., 2017; Constantino & Daw, 2015; Hayden et al., 2011; Kane et al., 2017; Le Heron et al., 2020). This assumption might not always be true (Merkle et al., 2017, 2014; Passingham, 1985; Riotte-Lambert et al., 2015; Sayers & Menzel, 2012), complicating optimal foraging decisions. The present study aimed to investigate patch-leaving decisions in an environment in which the same set of three patches could be revisited, and patches replenished their rewards at different rates. Within this environment, we manipulated whether participants had control over patch encounters, examining how this influenced decisions to leave a patch and which patches participants chose to visit next.

Consistent with our predictions (Hall-McMaster & Luyckx, 2019), we found that when participants were able to select which patch to exploit after a leave decision, they visited fast replenishing patches more often than slower replenishing patches. This resulted in higher reward rates, higher rewards when arriving at patches and higher reward outcomes prior to leave decisions, compared to environments in which patch encounters were randomly determined. We also predicted that participants would leave patches after fewer exploit actions in free choice blocks. However, increased rewards at arrival and prior to leaving patches in the free choice environment resulted in a comparable number of actions before leaving patches to the forced choice environment. To account for the behavioural effects above, we developed a simple logistic regression model for exploit-or-leave decisions, which included the previous reward outcome, the global reward rate across all patches, and the maximum reward rate among the alternative patches. We found that choices in free choice blocks were better explained with this model than an MVT-based model that included only the previous reward and the global reward rate. Two additional models that tested whether participants internally tracked patch replenishment did not provide a better description of the data. Despite behavioural differences between choice conditions, qualitatively similar results were found when modelling the forced choice blocks.

The present results build on previous studies based on Optimal Foraging Theory (Charnov, 1976; Stephens & Krebs, 1986), which involved random or pseudo-random encounters with new options when deciding to leave a current option (Barack et al., 2017; Constantino & Daw, 2015; Hayden et al., 2011; Kane et al., 2017; Le Heron et al., 2020), as well as theoretical studies examining patch-revisitation (Kilpatrick et al., 2020; Possingham & Houston, 1990). We show that allowing decision makers to select which options are pursued alters foraging behaviour. More frequent visits to high value options and less frequent visits to low value options increase the reward accrued per action, reflecting behaviour that is more efficient than encountering options at random and could improve biological fitness in natural settings (Merkle et al., 2017, 2014; Riotte-Lambert et al., 2015; Sayers & Menzel, 2012). The fact that exploit-or-leave choices were better explained with a model that used both global and local reward information suggests that humans do indeed use reward information about alternative patches to guide their exploit-or-leave decisions, in at least some foraging environments.

One interesting aspect of the results was that local reward rate information was included in the winning model for both free and forced choice conditions. One possible reason could be that, in the forced choice condition, participants still knew they would go to one of the two alternative patches. In this case, it could make sense to consider the mean reward rate across alternatives because the exact alternative that would be visited next was unknown. In the free choice condition, participants could have considered the maximum reward rate among the alternative patches because they were able to go directly to the desired patch after a leave decision. We did not have sufficient variability in candidate regressors to test this possibility. The maximum reward rate among the alternative patches and the mean across them were very highly correlated (r=0.92) and therefore, we only included the maximum alternative reward rate in our regression models. This means that participants could have been using different information about the alternative patches in the different choice conditions, but that it was not possible to detect this in the current setup.

Before running our main analyses, we excluded 10/70 participants who were outliers based on the total number of points earned (see STAR Methods). When looking in-depth, we found that these participants earned substantially fewer points due to fixed behavioural strategies, such as exploiting each patch until it was completely empty or leaving each patch after a single exploit action. While we took these stereotyped responses as evidence that these participants were not engaged with the task, it is possible that these behaviours reflect genuine strategies. To test whether excluding these participants influenced the results, we re-ran our statistical analyses with all 70 participants. This produced two changes in the results, neither of which undermined our main conclusions. First, the greater proportion of choices to the medium patch, observed when leaving the fast patch in the free choice blocks, was no longer significant (mean free choice=0.556, SD=0.199; mean forced choice=0.504, SD=0.130; corrected *p*=0.051; original result in Figure 2D: mean free choice=0.570, SD=0.189; mean forced choice=0.510, SD=0.112; corrected *p*=0.036). Second, the comparable number of exploit actions made before leaving the medium replenishing patch was now significantly lower in the free compared to the forced choice blocks (mean free choice=5.581, SD free choice=2.660 mean forced choice=5.995, SD forced choice=2.912; *t*(69)=2.931, corrected *p*=0.014; original result in Figure 4B: mean free choice=5.309, SD free choice=1.850; mean forced choice=5.581, SD forced choice=1.708; *t*(59)=2.207, corrected *p*=0.094).This contributed to a significantly fewer exploit actions before leaving overall, in the free choice blocks (mean free=5.531, SD=2.631; mean forced=5.869, SD=2.816; *F*(1,69)=6.652, *p*=0.012; original result: mean free=5.251, SD=1.772; mean forced=5.452, SD=1.643; *F*(1,59)=3.337, *p*=0.073). This could suggest that our main analysis with 60 participants did not have sufficient power to detect the leaving time effect. The effect size with 70 participants was small, and our impression is that future studies could use a more sensitive measure of leaving time (such as seconds in patch), or simulate agents in different task settings, to better understand how patch control influences leaving time. Altogether, the results with all 70 participants do not challenge the main conclusions. Even when all participants were included, control over patch encounters resulted in a higher proportion of visits to fast replenishing patches, patches being left at higher levels of reward patches, and higher average reward rates. In addition, decisions to exploit-or-leave a patch were still influenced by reward information about the alternative patches.

The main results have three broader connections to decision making research. In MVT, the decision maker needs to compare the instantaneous reward from the current patch with a long-run environmental reward rate. The latter quantity has been argued as an important variable for other kinds of choices, including the vigour of responding (Niv, 2007; Niv et al., 2007; Yoon et al., 2018), and there are suggestions (Niv, 2007; Niv et al., 2007) with at least some evidence (Hamid et al., 2016), that this variable is encoded in tonic concentrations of dopamine. One interesting characteristic of our foraging problem in this regard is that the relationship between long-run past reward rate and the immediate future reward is disrupted. Since different patches replenish at different rates in our task, the future reward rate is influenced by the replenishment status of the alternative patches. It is consequentially an interesting empirical question as to whether the leaving policy in foraging environments that allow patch revisiting might be implemented through changes in tonic dopamine or whether the neural mechanism for patch leaving might be quite different. In addition to the future rewards associated with replenishment, it could also be that participants are sensitive to accelerating reward depletion in the current patch, adapting their local learning rule for different depletion phases. Future studies could examine this by manipulating patch decay dynamics. A further consideration arises in the work of Garrett & Daw (2020), who considered a structurally different foraging problem, in which human participants could use a recent average reward rate the track the opportunity cost of time. The authors observed that participants used different learning rates for the opportunity cost, depending on whether rewards were better or worse than the recent average. While patch replenishment makes the present task more dynamic than the one in Garrett & Daw (2020), it would be interesting for future research to consider whether participants adopt different learning rates in relation to patch specific reward rates. Finally, the present work also explored a foraging problem that involved a small number of patches with structured replenishment and the possibility of revisitation. This is very different from the conventional case in which patches are too numerous to individuate. Indeed, it is closer to a problem class known as concurrent variable interval schedule with hold (Staddon et al., 1981). Nevertheless, it is interesting to consider intermediate cases, for instance in which there are very many patches, but they are systematically clustered in space according to their replenishment rates. This could allow animals to represent each individual patch in terms of the cluster it belongs to, and form a corresponding task representation (Schuck et al., 2016; Wilson et al., 2014; see Behrens et al., 2018; Niv, 2019 for reviews). This task representation could then be used to make decisions about when to spend time in which part of space. If there were additional sensory features that allowed distal prediction about the current state of a patch, animals could also readily learn about and exploit this information to act more efficiently.

Overall, the results from this experiment suggest that revisiting patches, and having control over when to do so, changes human foraging behaviour. In environments that involve patch-revisiting, leave decisions depend on the average reward for the environment, but also on the reward for alternative patches that might be revisited.

## Limitations of the Study

The present experiment had three main limitations. First, the experiment did not provide a computational account for how patches were selected. Participants tended to select patches with higher expected rewards when leaving all three patches and higher reward rate patches when leaving the medium replenishing patch. The winning model assumed that participants used the highest reward rate among the alternative patches to guide leaving behaviour, however, decision makers might in fact switch between different sources alternative patch information depending on the patch they are planning to select. Future efforts could therefore model both patch selection decisions and exploit-leave decisions within a unified framework. Second, participants did not know the replenishment speed for each island at the start of a block or the precise rates of replenishment. Behaviour at the beginning of a block might therefore reflect directed exploration, which is not accounted for in the models we present. Future work could therefore investigate how replenishment rate information is learned and investigate behavioural markers for shifting from learning about the environment to a more stable patch-exploitation strategy. Third, the experiment could be limited in its trial-based structure. While some patch-leaving designs are trial-based (Barack et al., 2017; Constantino & Daw, 2015; Kane et al., 2017; Hayden et al., 2011), studies tend to compute reward rates (Kane et al., 2017; Lottem et al., 2018), the probability of leaving (Constantino & Daw, 2015) or leaving times (Barack et al., 2017; Hayden et al., 2011; Le Heron et al., 2020; Lottem et al., 2018) using some continuous time information, such as the number of seconds in a patch. In the present experiment, we used trial-wise data to model discrete choice behaviour using a similar approach to Constantino & Daw, 2015. Unlike Constantino and Daw’s (2015) task, however, travel times were fixed in the present experiment and did not reduce the number of actions available in a block. Decisions about which patch to visit did count towards the number of actions in a block and travel time here refers to a 1s delay between patch selection and arriving at a patch. This meant we did not include temporal variables for exploit time and travel time in seconds within our models. While appropriate for our task design, this could have resulted in less sensitive reward rates and leaving time estimates, as compared with a continuous time-based setup.

## STAR Methods

### Resource Availability

#### Lead Contact

Further information and requests for resources and reagents should be directed to and will be fulfilled by the lead contact, Sam Hall-McMaster.

#### Materials Availability

This study used existing materials. Stimuli were presented using Psychopy-3 (https://www.psychopy.org/; similar to the version described in Peirce et al., 2019). Stimuli included three circles, which were used as different choice options, a cartoon pirate ship, and a cartoon medallion, used for reward feedback (see Figure 1). Cartoon stimuli were created by icon developers Smalllikeart and Nikita Golubev, and accessed at https://www.flaticon.com. Stimulus sizes were scaled based on participants’ screen sizes and responses were recorded using participants’ keyboards. Participants were asked to only take part if their screen was at least 13 inches and they were using Chrome as their browser. The task was hosted on Pavlovia (https://pavlovia.org/) which stored participant data during online testing. Data were analysed using MATLAB (RRID: SCR_001622, version 2019b) and RStudio (RRID: SCR_000432, version 1.4.1717).

**Figure 1.**
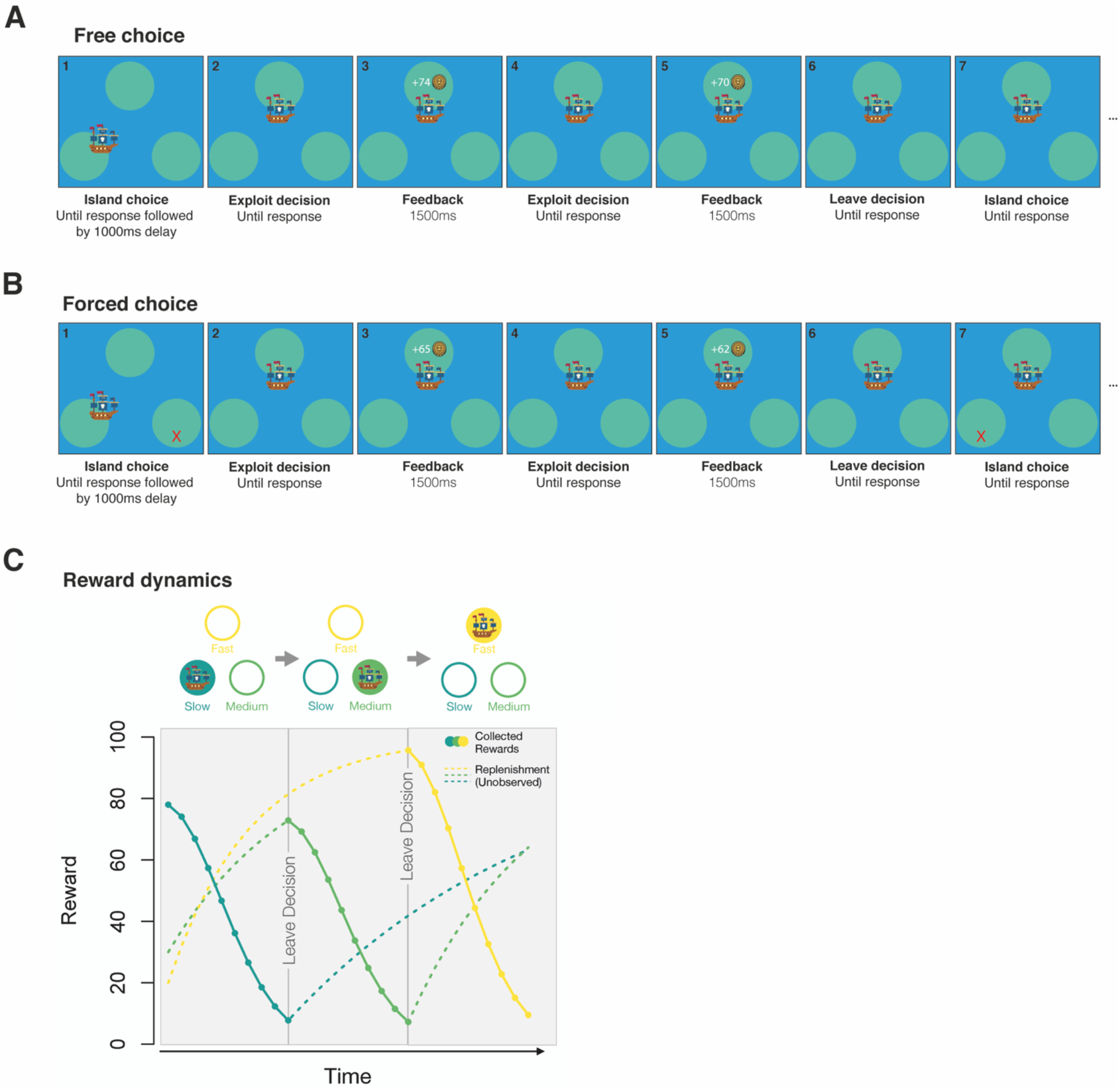
Example trial sequences for free and forced choice conditions. A: Free choice trials. Participants first chose an island to sail to (screen 1). Once their ship had arrived, participants could dig on the island for buried treasure (an exploit action shown on screen 2). Following a dig, participants received feedback about the number of coins added to their treasure chest (screen 3). Participants could continue digging (screens 4-5), until deciding to leave the island (screen 6) and choosing another island to sail to (screen 7). B: Forced choice trials. Under forced choice conditions, the seas were stormy and islands were not always accessible. Participants were forced to choose just one of the two alternative islands, after a leave decision. The accessible island was randomly determined after each leave decision and the other, inaccessible island was marked with a red X. All other aspects were as in the free choice condition shown in panel A. C: Reward dynamics. When an island was being exploited for treasure, the number of coins added to the treasure chest decreased with each successive dig. The depletion process during exploitation was the same for all islands. While an island was being exploited, new coins were buried on the alternative island at island-specific rates (slow, medium or fast, shown in sea green, green and yellow lines respectively). Participants did not see new coins being buried and therefore needed to revisit islands to learn about their replenishment speeds. No reward was given for choosing an island to sail to or deciding to leave an island (not indicated in panel).

#### Data and Code Availability

Data, task and analysis code are publicly available on the Open Science Framework (DOI: 10.17605/OSF.IO/C4TRJ; https://osf.io/c4trj/). Any additional information required to reanalyze the data reported in this paper is available from the lead contact upon request.

### Experimental Model and Subject Details

70 human participants (mean age=29, 38 female, 32 male) took part in this experiment as an online study, which ran using Prolific (https://www.prolific.co/). Participants were eligible to take part if they were 18-40 years of age, fluent in English and were not receiving treatment or taking medication for mental illness. Eligible participants were screened with attention checks before participation was allowed. Ten participants, from the 70 who passed the attention checks and completed the experiment, were excluded due to their task performance (number of points earned) falling more than three median absolute deviations below the sample median. For six rejected participants, this low performance was due to a fixed response profile, in which participants exploited each patch until it was empty before leaving. This response profile been used as a basis for exclusion in previous human foraging studies (Constantino & Daw, 2016; Lenow et al., 2017). Four rejected participants showed low performance due to the opposite strategy, in which a patch was often exploited just once before leaving. These fixed response strategies suggest that the ten participants marked for rejection did not make engaged foraging choices during the task. Excluding these participants did not affect our main conclusions (see Discussion for additional details). The remaining 60 participants were between 18 and 40 (mean age=28, 33 female, 27 male). We had no reason to expect, nor was there evidence for, an effect of gender on performance (mean points earned female=3.758e4, SD=2.134e3; mean points earned male=3.778e4, SD=1.864e3; unpaired *t*(58)=0.399, *p*=0.692). Participants received £5 per hour and could earn up to £1 extra for depending on their task performance. The study was approved by the Max Planck Institute for Human Development Ethics Committee (N-2020-06) and all participants indicated their informed consent before taking part.

### Method Details

Participants performed a patch-leaving task, based on principles from Optimal Foraging Theory (Stephens & Krebs, 1986). The classic patch-leaving setup involves participants exploiting an option as the reward it returns decreases over time. Participants’ central choice is when the option is no longer worth exploiting; when it should be abandoned to search for something better. Once participants decide to leave the option, a new option is presented for them to exploit, the value of which is independent of their previous actions. We made three critical adaptations to this standard setup. First, we manipulated whether participants could choose their next option, after a leave decision. Second, participants revisited the same three options within a block, which meant that their previous decisions would influence an option’s reward value when returning to it. Third, each option replenished its reward over time, but at a different rate.

Participants played the role of a pirate, sailing to different islands in search of buried treasure (Figure 1). The common elements across all conditions of the task were as follows. Participants first selected an island to sail to using the keys F (left island), J (upper island) and K (right island). Following a travel delay of 1000ms to reach the island, participants made a series of exploit-leave decisions. When deciding to exploit the island for treasure (space bar press), participants received between 0 and 100 gold coins, which were indicated on screen for 1500ms. The reward feedback then disappeared and participants made another exploit-leave decision. This loop continued until the participant decided to leave the island (S key press), following which they were able to select a new island to sail, and thereafter enter a new exploit-leave loop. Participants needed to perform at least one exploit action before being able to leave an island. The event sequence described above continued until the participant had performed a total of 200 actions, at which point the block ended. Choices about which island to sail to, exploit decisions and leave decisions all counted as individual actions. The actions remaining in a block were shown in the top right corner of the screen. While sailing between islands took time, reducing the reward per unit time, it did not affect the reward per action because sailing did not consume the actions available in a block. We therefore used the reward per action as our main measure of reward rate. The critical manipulation between blocks was whether participants had a free or forced choice over which island they sailed to, after each leave decision. In free choice blocks, the seas were calm and participants could choose to visit either of the two alternative islands, after a leave decision. In forced choice blocks, the seas were stormy and islands were not always accessible. Participants were therefore forced to select one of the alternative islands, which was randomly determined at each selection phase. Inaccessible islands were marked with red X symbols.

The reward dynamics for each island functioned the same way across task conditions. When visiting an island/patch, *p*, the number of coins gained for the first exploit action was equal to the full number of coins buried. When a subsequent exploit action was made, the reward gained at time *t*, (denoted as *r*_*p,t*_) decreased following the equation:

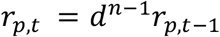

The decay constant, *d*, was set to 0.95, *n* refers the number of exploit actions since arriving at the island, and *r*_*p,t*−1_ refers to the reward gained for the previous exploit action. Note that the decay constant declines exponentially in this expression, leading to accelerating decay characterized by a “squared exponent” relationship between the reward when entering the patch at time *t* and the reward after making *k* exploit actions prior to the current exploit action (i.e. *k* = *n*-1):

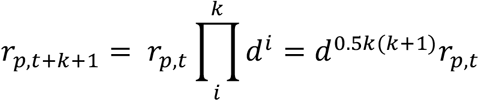

While one island was being exploited, new coins were buried on the other two islands. The coins buried on each alternative island *q* increased, following each decision to exploit the current island, according to the equation:

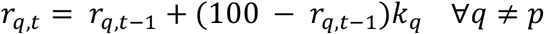

The variable *k*_*q*_ is the replenish rate for alternative patch *q*. The replenish rate was different for each island, with one island replenishing at a slow rate (0.05), one at a moderate rate (0.10) and one at a fast rate (0.15). This equation was also used to update the coins buried on all islands, after island selection responses and leave responses. Each island could replenish to a maximum of 100 coins. In each block, the slow, medium and fast replenishing island was randomly assigned to the left, top or right position on screen.

Participants completed four blocks, two free choice blocks and two forced choice blocks. The block order was random. Participants were informed that each block would contain a fast, slow and medium replenishing island but did not have direct experience with the islands before starting the experiment. Participants were instructed about whether an upcoming block would be free or forced choice and were reminded to gain as much reward as possible. The block began with a choice between all islands (free choice) or a randomly selected island (forced choice). Islands were located at one of three vertex positions on screen that formed an equilateral triangle. However, slow, medium and fast replenishing islands were randomly assigned to the vertex positions in each block. Participants therefore needed to revisit different islands (left, top, right) to learn about their replenishment speeds when starting a new block. The coins buried at each island were initialised to random numbers between 69 and 79 at the beginning of the block. This range was determined based on simulating 1000 agents that completed one forced choice block, using the average experienced reward rate to make leave decisions. In these simulations, the initial reward for each option was set to 100. Average experienced reward rates were calculated prior to each stay-leave decision, by dividing the total reward gained by the total number of actions taken. The reward gained for the first exploit action when arriving at an island was recorded during the second half of the block, allowing time for the islands to reach steady reward dynamics. We then calculated 95% confidence intervals for the recorded values and averaged confidence intervals across agents, resulting in a lower bound of 69.3521 and an upper bound of 79.1659 points.

### Quantification and Statistical Analysis

#### Visitation

To test the prediction that participants would revisit high value options more and low value options less under free choice conditions, we extracted the number of times each patch (slow, medium, fast replenishment rate) was selected in forced and free choice blocks. The number for each patch in each condition was then normalised by the total number of selection actions performed in its respective condition. For example, the number of visits to the fast patch in the free choice blocks was divided by the total number of visits made to any patch in the free choice blocks. This normalisation step means that the proportion of visits to each patch in the forced choice condition sum to one and the proportion of visits to each patch in the free choice condition sum to one. Normalised visits under free and forced choice conditions were compared for each patch using separate paired two-tailed t-tests. In exploratory analyses, we examined how often the alternative patch with the highest expected reward (number of coins buried) was chosen, when leaving each patch. For each patch being left (slow, medium, fast replenishment rate), we extracted the number of coins for each alternative patch at the time of choosing the next patch to sail to. We then computed the proportion of times participants chose the alternative patch with the highest number of coins. Proportions for each patch being left were compared between choices using separate two-tailed t-tests.

To address potential issues with non-normally distributed (bounded) proportions, we used non-parametric permutation testing when assessing statistical differences in all tests involving proportions. This approach was selected because it did not assume a specific data distribution (e.g. Gaussian). When testing two proportions (e.g. the proportion of choices to the best alternative for free versus forced choice), the permutation procedure randomly swapped the condition data for about half the participants. This meant that, for approximately 50% of the participants, the forced choice proportion became the free choice proportion and vice versa. The procedure then conducted a paired t-test and stored the absolute value of the resulting *t*-statistic. This was repeated 10,000 times to generate a null distribution of *t*-values for a given statistical test. The absolute *t*-value from the true *t*-test was then compared against the 95th percentile of the null distribution. If it was higher, the test was significant at an alpha level of 0.05. The corrected *p*-value was computed based on the percentile of the true *t*-value within the null distribution. In cases where the true *t*-value was higher than all values in the null distribution, the *p*-value is one over the number of permutations (i.e. 1e-4). The permutation procedure was implemented with custom MATLAB code and verified with an independent toolbox for permutation testing (https://www.mathworks.com/matlabcentral/fileexchange/29782-mult_comp_perm_t1-data-n_perm-tail-alpha_level-mu-reports-seed_state), which produced the same results.

#### Average Reward Rates

To test the prediction that the average reward rate would be higher under free choice conditions, we extracted the number of points earned in forced and free choice blocks. These values were then divided by the number of exploit actions in each condition. Once this procedure had been performed for each participant, reward rates were compared using a 2 × 3 repeated measures ANOVA, with factors of choice condition (free × forced) and patch (slow, medium, fast replenishment rate). The interaction between choice condition and patch type was followed up with paired t-tests that compared whether the difference in reward rate between choice conditions (i.e. free minus forced) differed between patch pairs (slow vs medium, slow vs fast, medium vs fast). We conducted additional t-tests to examine whether the increase in reward rate under free choice conditions was significantly different from zero for each patch (as opposed to the interaction arising from significant reward rate increases for some patches but not others). The alpha threshold for each set of t-tests was corrected for three exploratory tests using the Bonferroni correction. Reward rates based on all actions (including exploiting, leaving and patch selection) were also calculated and are reported in the main text.

#### Reward Thresholds for Leaving

To test the prediction that participant’s reward thresholds for leaving the current patch would be higher in free choice blocks, we extracted the last reward outcome prior to each leave decision. We then averaged the reward outcome before leaving, separately for each choice condition. Once this procedure had been performed for all participants, a 2 × 3 repeated measures ANOVA was performed on the reward outcome before leaving, with factors of choice condition (free × forced) and patch (slow, medium, fast replenishment rate). The main effect of patch was followed up with paired t-tests that compared the mean reward before leaving between the fast and medium replenishing patches, the fast and slow replenishing patches, as well as the medium and slow replenishing patches. The alpha threshold was corrected for three follow up tests using the Bonferroni correction. We also note here that the term leaving threshold was used in a slightly different way in Hall-McMaster and Luyckx (2019). In that paper, the leaving threshold was used to refer to the effort/time invested in a patch before leaving. We therefore wrote that participants would have a lower threshold for leaving under free choice, meaning that less time would be spent exploiting a patch and it would be left at a higher level of reward. In this paper, we used leaving threshold to mean the reward gained from the current patch, immediately prior to a leave decision. When used in this way, the same prediction is expressed as the free choice condition producing a higher leaving threshold (i.e. a patch is left at a higher level of reward). The prediction is the same in both cases but we hope this explanation of how the terms were used helps to allay any confusion readers could have when reading across the two papers.

#### Rewards on Arrival

To examine rewards when participants arrived at patches, we extracted the number of coins earned for the first exploit action, each time the slow, medium and fast replenishing patches were visited. We then averaged the arrival rewards, for each patch and each condition. We then performed a 2×3 repeated measures ANOVA on the arrival rewards, with factors of choice condition (free × forced) and the arrival patch (slow, medium, fast replenishment rate). The interaction was followed up with paired t-tests that compared the arrival reward between choice conditions, separately for each arrival patch. The alpha threshold for the three exploratory tests was corrected using the Bonferroni correction. A similar procedure was performed to examine the average arrival reward, as a function of the patch being left. The critical difference was that we extracted the first reward when arriving at the next patch (regardless of its replenishment rate), each time participants decided to leave the slow, medium or fast replenishing patch. The main effect of the patch being abandoned was followed up with separate paired t-tests that compared mean arrival rewards between specific patches (i.e. slow vs medium, slow vs fast, medium vs fast). The alpha threshold was therefore corrected for three exploratory tests using the Bonferroni correction

#### Actions Before Leaving

To test the prediction that participants would make fewer exploit actions prior to leaving patches under free choice conditions, we extracted the number of exploit actions before each leave decision. We then averaged the exploit actions, separately for each choice condition. Once this procedure had been performed for all participants, a 2 × 3 repeated measures ANOVA was performed on the number of exploit actions before leaving, with factors of choice condition (free × forced) and patch (slow, medium, fast replenishment rate). In addition, we performed exploratory analyses that examined whether leaving times differed between choice conditions for each patch individually. Exploratory analyses followed the same procedure above, except that the number of exploit actions prior to leaving were averaged separately for each patch and compared between choice conditions using separate paired two-tailed t-tests. The alpha threshold for the three exploratory tests was corrected using the Bonferroni correction.

#### Simulated Performance

We first used dynamic programming to determine the optimal policies for a slightly simpler version of both free and forced tasks, and then used a form of certainty equivalent control based on inferences about the states of the patches in order to choose actions. We expect the approximations inherent in the simpler tasks to be relatively benign. The simpler versions of the task assumed that the agent always had full knowledge of which patch was which (including how fast each one replenished), the current value of each patch, and rounded all these values to the nearest integer (consistent with the integer rewards provided in the task). The agent also knew how many actions were remaining in the block. We then used the Bellman equation, working backwards from the last of the 200 actions in a block to the first to find the optimal policies for the state of the patches and the agent in terms of whether to leave the current patch, and, if so, which patch to choose next (when this choice is possible during the free choice blocks). The value of an action when there are n actions remaining depends on the values after there are only n-1 actions remaining, which is why working backwards is appropriate. These policies end up being very high dimensional, since there are 200 actions per block, each of the patches has 101 possible states, and because the depletion is not memoryless (i.e. the rate of depletion depends on the number of timesteps the agent has spent in a patch), the policies also vary as a function of current patch identity and the time spent in it.

Given these policies, the agent is then placed in the same environment as the subjects, in that it only actually knows the distribution of the starting values of the patches, and does not know which one is which. It therefore performs Bayesian inference to estimate the identity of the patches. In making choices, one approximation it makes is to assume the maximum likelihood assignment of these identities (this is almost perfect after two visits, given the difference in replenishment rates). A second approximation is that the agent assumes all patches start off at the same value. We searched to find the starting value assumed by the agent that optimized overall reward. Assumed starting values between 80-95 did not influence the average reward accrued by agents in the free choice blocks, presumably because these values are rapidly corrected. In forced choice blocks, the variability in average reward due to random patch selection had a larger influence on average reward than variation in starting value. As it did not affect the results, we used 95 as the assumed starting value in the simulations. We generated 1000 agents performing two free choice and two forced choice blocks. Mean scores across agents are shown on Figures 2-4.

**Figure 2.**
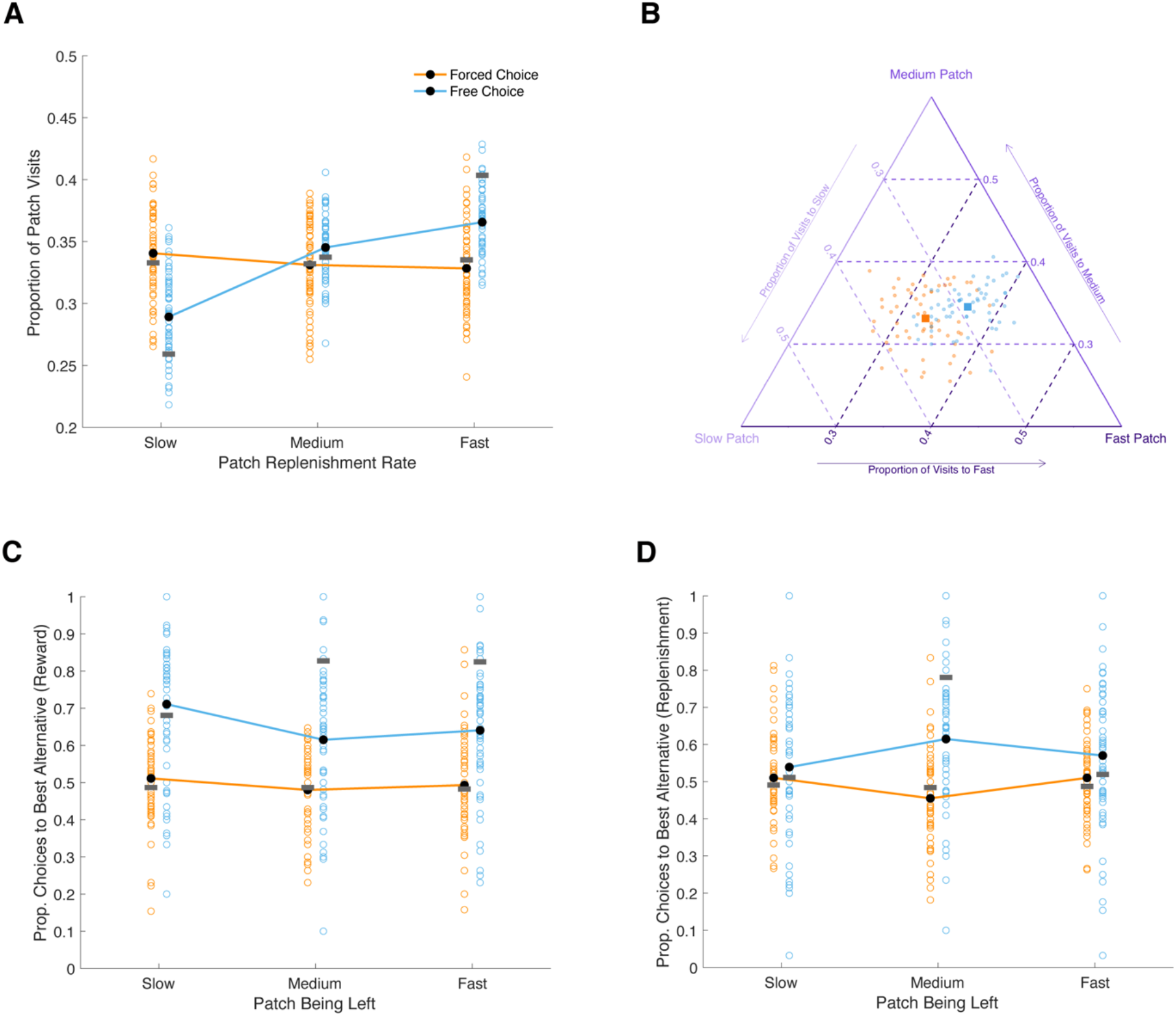
Patch choices. A: Proportion of visits to the slow, medium and fast replenishing patch, under forced and free choice conditions. Orange colours indicate data from forced choice blocks and blue colours indicate data from free choice blocks. Unfilled circles show proportions for individual participants and lines connect average proportions across the sample. Horizontal grey bars overlaid on each condition show the mean performance of simulated agents that approximately maximise reward on the task. Proportions were could deviate from 0.33 in the forced choice condition for individual participants because the next patch was selected at random (with a probability of 0.5) after each leave decision. B: The same data as panel A, but plotted as a single point on a simplex for each participant. Squares indicate mean visit proportions for each choice condition. For A and B, proportions across the three patches sum to one within each choice condition. C: Proportion of choices to the alternative patch with the higher expected reward (i.e. current patch value), when leaving the slow, medium and fast replenishing patches. D: Proportion of choices to the alternative patch with higher replenishment, when leaving the slow, medium and fast patches. For B, C and D, colours are the same as those used in panel A.

**Figure 3.**
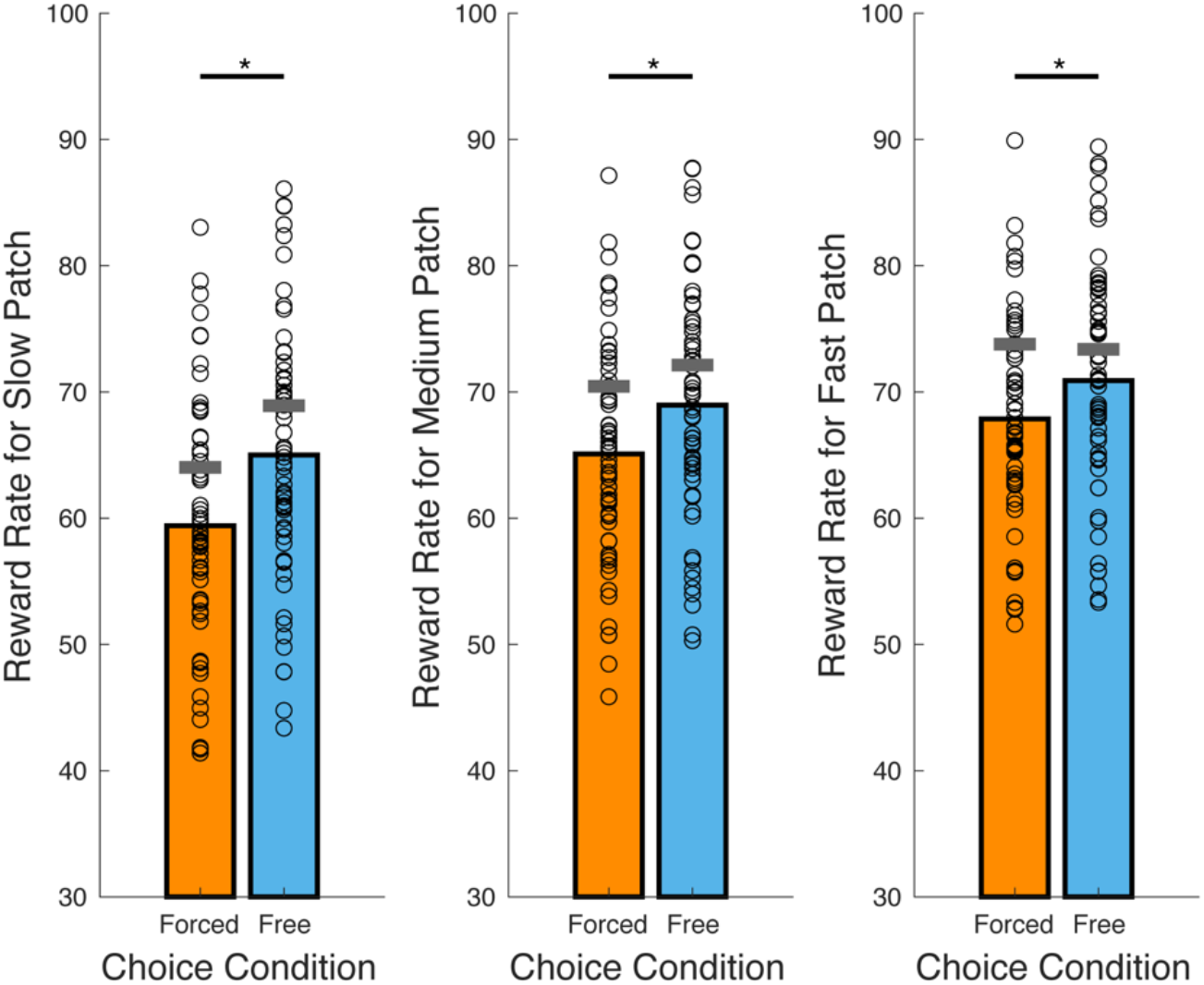
Reward per exploit action (reward rate) under free and forced choice conditions, in the slow, medium and fast replenishing patch. Coloured bars show mean reward rates for each choice condition. Circles overlaid on each bar show individual participant values. Horizontal grey bars overlaid on each condition show the mean performance of simulated agents that approximately maximise reward on the task. *p<0.05.

**Figure 4.**
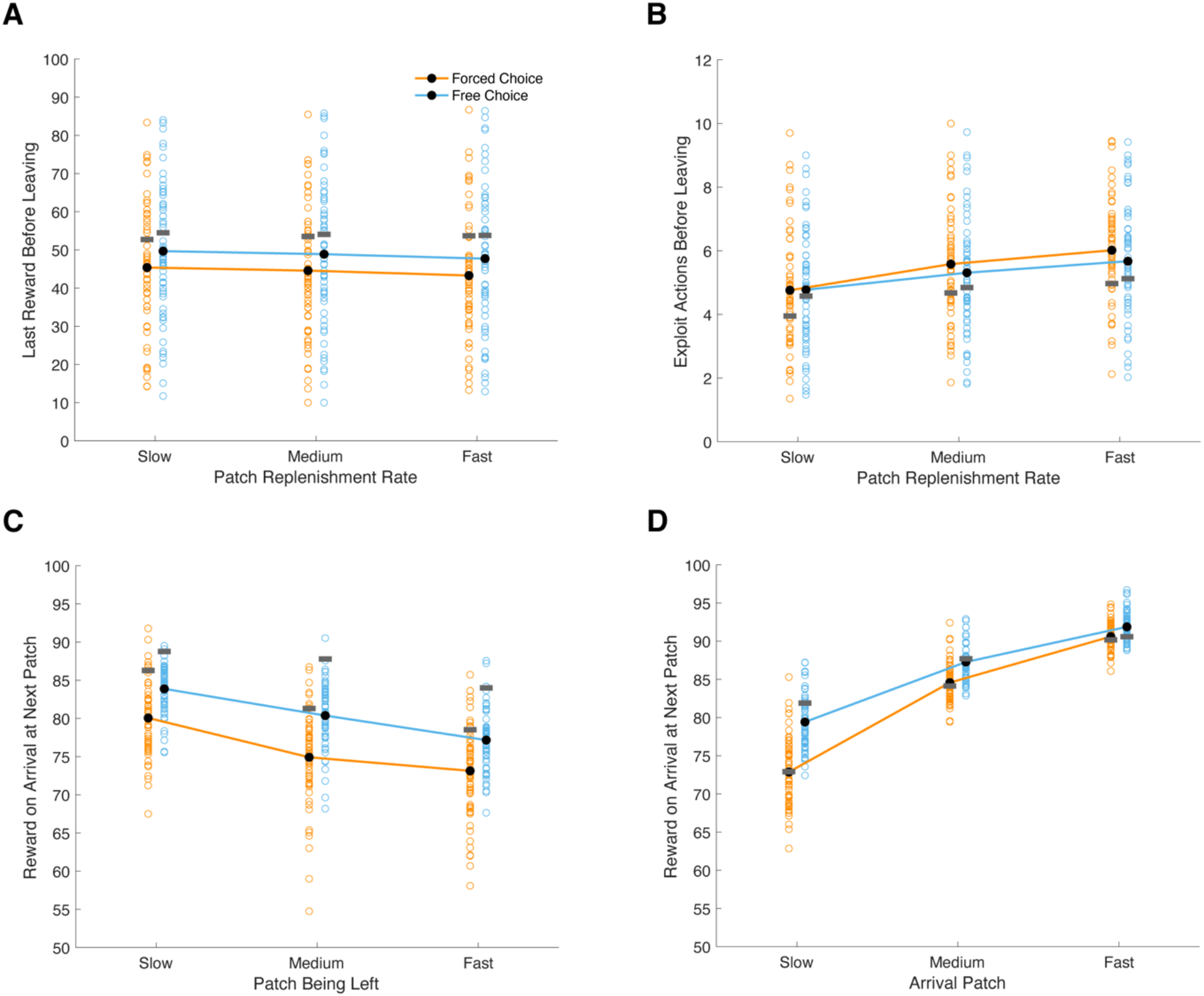
A-B: Patch-leaving characteristics. A: Last reward prior to leaving the slow, medium and fast replenishing patch. Orange indicates data from forced choice blocks and blue indicates data from free choice blocks. Black dots indicate averages and unfilled circles show values for individual participants. Horizontal grey bars overlaid on each condition show the mean values for simulated agents that approximately maximise reward on the task. B: The number of exploit actions made before deciding to leave each patch. C-D: Patch-arrival characteristics. C: Average reward for the first exploit action at a new patch, after leaving the slow, medium or fast replenishing patch. D: Average reward for the first exploit action when arriving at the slow, medium or fast replenishing patch. For B, C and D, colours are the same as those used in panel A.

**Figure 5.**
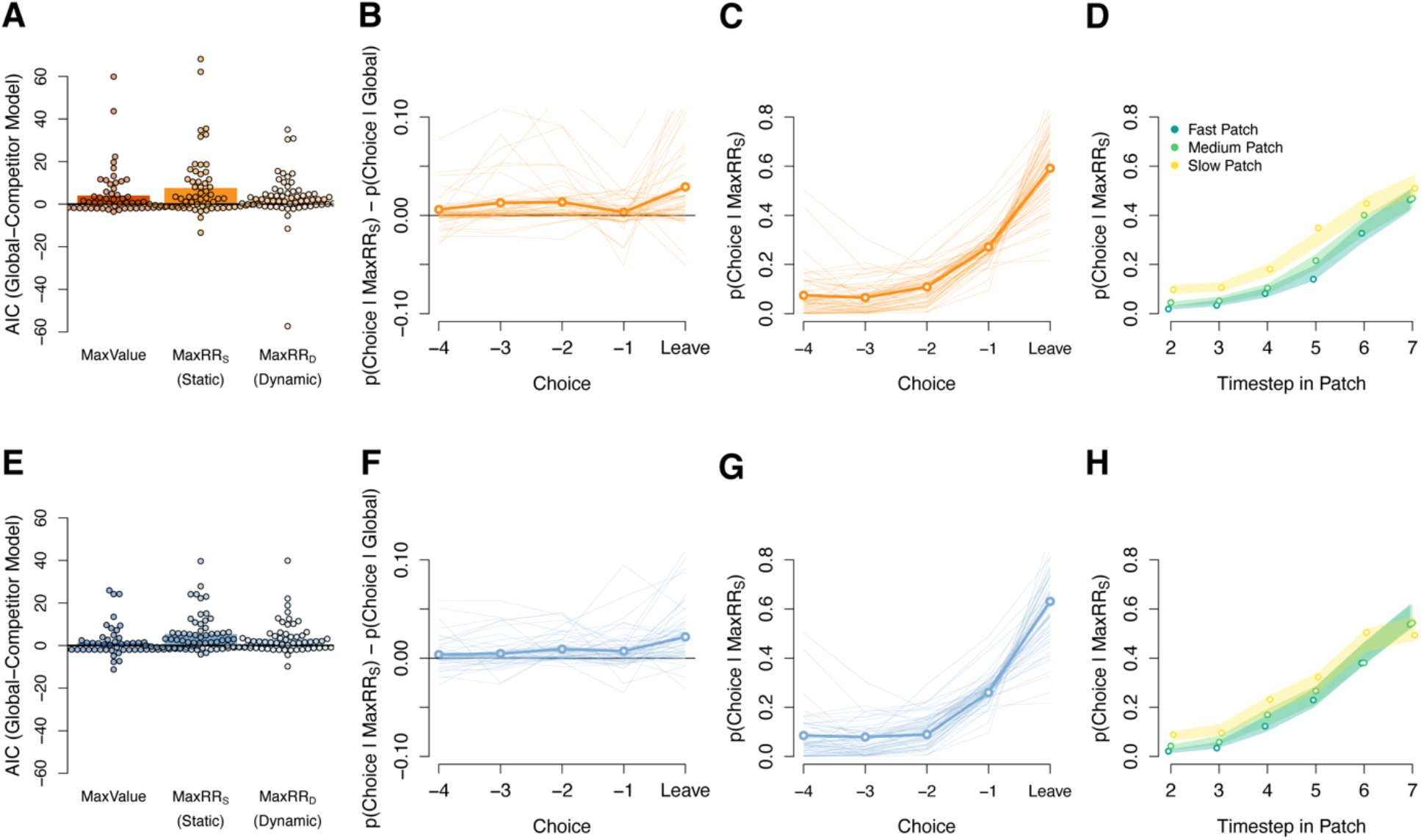
Logistic regression models for exploit-or-leave decisions. A: Differences in AIC scores between the global model (used as a baseline) and each *global+* competitor model described in the text, when fit to forced choice blocks. Bars indicate the group means and dots represent individual participant data. A positive difference score indicates greater evidence for the *global+* model over the global model. The most consistent improvement was achieved with the *global+maxRR*_*S*_ model, which included the maximum estimated reward rate among the alternative patches as a predictor for patch-leaving. B: The timecourse of choice probabilities for the winning *global+maxRR*_*S*_ model, relative to the global model. Positive numbers indicate a closer match to choice behaviour under the *global+maxRR*_*S*_ model. Probabilities are plotted as a function of timestep in the patch, in the leadup to a leave decision. On average, the winning model maintains a numerical advantage for exploit decisions (first 4 dots) as well as leave decisions (rightmost dot). Thin lines show data for individual participants and the bold line shows the sample mean across. Data are averaged over all patch types. C: The absolute probability of a leave decision under the *global+maxRR*_*S*_ model. Dots, thin lines and bold lines have the same meaning as panel B. D: The probability of leaving a patch as a function of timesteps in the current patch, separated for slow, medium and fast replenishing patches. Dots show the probabilities for participants’ actual choices. Shaded lines show the corresponding probabilities of leaving under the *global+maxRR*_*S*_ model, with the shading width showing standard errors of the mean. The colours corresponding to each patch are shown in the panel legend. Panels E-H are analogous to panels A-D, but with data from the free choice condition.

#### Use of Global and Local Reward for Leave Decisions

Our main analysis concerned whether exploit-or-leave choices could be better predicted using global and local reward information, rather than global reward information alone. To investigate this, we conducted a series of logistic regressions. First, we constructed a global regression model that predicted exploit-or-leave choices using the reward gained from the previous action and the average reward rate. Three free parameters were used to estimate the average reward rate on each trial before running the logistic regression. The average reward rate was initalised at a starting value, *s*, which was constrained between +50 and +100. The average reward rate estimate was updated after each exploit action using a simple delta learning rule (learning rate *α*) and after each leave action (learning rate *α*_*l*_). After calculating the average reward rates for each trial, we removed the first exploit action when arriving at a patch from the analysis. This was done since participants had to perform at least one exploit action when arriving at a new patch and, therefore, these trials cannot be considered true exploit-or-leave choices. We also excluded trials from the last patch visit in each block, in which no leave decision was made. After the final data frame had been created, the logistic regression was performed. Each regression included a constant, plus the previous reward and average reward regressors. Model fitting was performed using a coordinate descent approach, whereby on each iteration the free parameters used to estimate the reward rate, *s, α* and *α*_*l*_, were fixed and the best coefficients of the logistic regression, one for the constant term, *β*_0_, and one for each predictor, *β*_*n*_, were determined using iteratively reweighted least squares. The likelihood of the data was then calculated and a new choice of the 3 hyper-parameters was determined using the function nloptr in R, which employed a DIviding RECTangles search algorithm.

Our modelling followed a nested model comparison approach, in which the global model served as a baseline and we added local task variables as predictors one by one. This allowed us to generate new (*global+*) models and test whether adding local information improved the fit to choice behavior over the baseline model. A description of each *global+* model is presented in the main text. Performance between each *global+* model and the global model was evaluated through comparison of the Akaike Information Criterion (AIC; Akaike, 1974) and the Bayesian Information Criterion (BIC; Schwarz, 1978). The three hyper-parameters used to estimate the reward rates (*s, α* and *α*_*l*_) were not included in the AIC and BIC calculations. However, we were primarily interested in the difference in AIC and BIC scores between models, which should still be accurate using this approach.

## Acknowledgments

This research was funded by an Alexander von Humboldt Foundation Fellowship awarded to S.H., funding from Max Planck Society and Alexander von Humboldt Foundation awarded to P.D., as well as an Independent Max Planck Research Group grant from the Max Planck Society (M.TN.A.BILD0004) and a Starting Grant from the European Union (ERC-2019-StG REPLAY-852669) awarded to N.W.S. The research also received financial support from the Max Planck Institute for Human Development. The authors thank Nir Moneta and Christoph Koch for technical advice, as well as Noa L. Hedrich for comments on the manuscript.

## Author Contributions

Conceptualisation, S.H., P.D., N.W.S; Methodology, S.H., P.D., N.W.S; Investigation, S.H.; Writing - Original Draft, S.H; Writing - Review and Editing, S.H., P.D., N.W.S; Visualisation, S.H, N.W.S.; Supervision, N.W.S; Funding Acquisition, N.W.S.

## References

Akaike, H. (1974). A new look at the statistical model identification. IEEE transactions on Automatic Control, 19(6), 716–723.

Barack, D. L., Chang, S. W. C., & Platt, M. L. (2017). Posterior cingulate neurons dynamically signal decisions to disengage during foraging. Neuron, 96(2), 339–347.

Behrens, T. E., Muller, T. H., Whittington, J. C., Mark, S., Baram, A. B., Stachenfeld, K. L., & Kurth-Nelson, Z. (2018). What is a cognitive map? Organizing knowledge for flexible behavior. Neuron, 100(2), 490–509.

Berger-Tal, O., & Avgar, T. (2012). The glass is half-full: overestimating the quality of a novel environment is advantageous. PLoS One, 7(4), e34578.

Charnov, E. L. (1976). Optimal foraging, the Marginal Value Theorem. Theoretical Population Biology, 9, 129–136.

Constantino, S. M., & Daw, N. D. (2015). Learning the opportunity cost of time in a patch-foraging task. Cognitive, Affective & Behavioral Neuroscience, 15(4), 837–853.

Erwin, R. M. (1985). Foraging decisions, patch use, and seasonality in egrets (Aves: Ciconiiformes). Ecology, 66(3), 837–844.

Fouragnan, E. F., Chau, B. K., Folloni, D., Kolling, N., Verhagen, L., Klein-Flügge, M., … & Rushworth, M. F. (2019). The macaque anterior cingulate cortex translates counterfactual choice value into actual behavioral change. Nature Neuroscience, 22(5), 797–808.

Garcia, L. C., & Eubanks, M. D. (2019). Overcompensation for insect herbivory: A review and meta-analysis of the evidence. Ecology, 100(3), e02585.

Garrett, N., & Daw, N. D. (2020). Biased belief updating and suboptimal choice in foraging decisions. Nature communications, 11(1), 1–12.

Gershman, S. J., Norman, K. A., & Niv, Y. (2015). Discovering latent causes in reinforcement learning. Current Opinion in Behavioral Sciences, 5, 43–50.

Hall-McMaster, S., & Luyckx, F. (2019). Revisiting foraging approaches in neuroscience. Cognitive, Affective, & Behavioral Neuroscience, 19(2), 225–230.

Hamid, A. A., Pettibone, J. R., Mabrouk, O. S., Hetrick, V. L., Schmidt, R., Vander Weele, C. M., … & Berke, J. D. (2016). Mesolimbic dopamine signals the value of work. Nature Neuroscience, 19(1), 117–126.

Hayden, B. Y., Pearson, J. M., & Platt, M. L. (2011). Neuronal basis of sequential foraging decisions in a patchy environment. Nature Neuroscience, 14, 933–939.

Huys, Q. J., Eshel, N., O’Nions, E., Sheridan, L., Dayan, P., & Roiser, J. P. (2012). Bonsai trees in your head: how the pavlovian system sculpts goal-directed choices by pruning decision trees. PLoS Comput Biol, 8(3), e1002410.

Huys, Q. J., Lally, N., Faulkner, P., Eshel, N., Seifritz, E., Gershman, S. J., … & Roiser, J. P. (2015). Interplay of approximate planning strategies. Proceedings of the National Academy of Sciences, 112(10), 3098–3103.

Kamil, A. C. (1978). Systematic Foraging by a Nectar-Feeding Bird, the Amakihi (Loxops virens). Journal of Comparative and Physiological Psychology, 92(3), 388–396.

Kane, G. A., Vazey, E. M., Wilson, R. C., Shenhav, A., Daw, N. D., Aston-Jones, G., & Cohen, J. D. (2017). Increased locus coeruleus tonic activity causes disengagement from a patch-foraging task. Cognitive, Affective and Behavioral Neuroscience, 17(6), 1–11.

Kilpatrick, Z. P., Davidson, J. D., & Hady, A. E. (2020). Normative theory of patch foraging decisions. arXiv preprint arXiv:2004.10671.

Kolling, N., Behrens, T. E. J., Mars, R. B., & Rushworth, M. F. S. (2012). Neural mechanisms of foraging. Science, 336(6077), 95–98.

Kolling, N., Wittmann, M. K., Behrens, T. E. J., Boorman, E. D., Mars, R. B., & Rushworth, M. F. S. (2016). Value, search, persistence and model updating in anterior cingulate cortex. Nature Neuroscience, 19(10), 1280–1285.

Kurth-Nelson, Z., Economides, M., Dolan, R. J., & Dayan, P. (2016). Fast sequences of non-spatial state representations in humans. Neuron, 91(1), 194–204.

Le Heron, C., Kolling, N., Plant, O., Kienast, A., Janska, R., Ang, Y. S., … & Apps, M. A. (2020). Dopamine modulates dynamic decision-making during foraging. Journal of Neuroscience.

Lenow, J. K., Constantino, S. M., Daw, N. D., & Phelps, E. A. (2017). Chronic and acute stress promote overexploitation in serial decision making. Journal of Neuroscience, 37(23), 5681–5689.

Lihoreau, M., Chittka, L., & Raine, N. E. (2010). Travel optimization by foraging bumblebees through readjustments of traplines after discovery of new feeding locations. The American Naturalist, 176(6), 744–757.

Lottem, E., Banerjee, D., Vertechi, P., Sarra, D., Lohuis, M. O., & Mainen, Z. F. (2018). Activation of serotonin neurons promotes active persistence in a probabilistic foraging task. Nature Communications, 9(1), 1–12.

Marshall, H. H., Carter, A. J., Ashford, A., Rowcliffe, J. M., & Cowlishaw, G. (2013). How do foragers decide when to leave a patch? A test of alternative models under natural and experimental conditions. Journal of Animal Ecology, 82(4), 894–902.

Merkle, J. A., Fortin, D., & Morales, J. M. (2014). A memory-based foraging tactic reveals an adaptive mechanism for restricted space use. Ecology Letters, 17(8), 924–931.

Merkle, J. A., Potts, J. R., & Fortin, D. (2017). Energy benefits and emergent space use patterns of an empirically parameterized model of memory-based patch selection. Oikos, 126(2), 185–195.

Mobbs, D., Trimmer, P. C., Blumstein, D. T., & Dayan, P. (2018). Foraging for foundations in decision neuroscience: Insights from ethology. Nature Reviews Neuroscience, 19, 419– 427.

Niv, Y. (2007). Cost, benefit, tonic, phasic: what do response rates tell us about dopamine and motivation? Annals of the New York Academy of Sciences, 1104(1), 357–376.

Niv, Y. (2019). Learning task-state representations. Nature Neuroscience, 22(10), 1544–1553.

Niv, Y., Daw, N. D., Joel, D., & Dayan, P. (2007). Tonic dopamine: opportunity costs and the control of response vigor. Psychopharmacology, 191(3), 507–520.

Passingham, R. E. (1985). Memory of monkeys (Macaco mulatta) with lesions in prefrontal cortex. Behavioral Neuroscience, 99(1), 3–21.

Peirce, J., Gray, J. R., Simpson, S., MacAskill, M., Höchenberger, R., Sogo, H., … & Lindeløv, J. K. (2019). PsychoPy2: Experiments in behavior made easy. Behavior Research Methods, 51(1), 195–203.

Possingham, H. P., & Houston, A. I. (1990). Optimal patch use by a territorial forager. Journal of Theoretical Biology, 145(3), 343–353.

Riotte-Lambert, L., Benhamou, S., & Chamaillé-Jammes, S. (2015). How memory-based movement leads to nonterritorial spatial segregation. The American Naturalist, 185(4), E103–E116.

Rushworth, M. F., Noonan, M. P., Boorman, E. D., Walton, M. E., & Behrens, T. E. (2011). Frontal cortex and reward-guided learning and decision-making. Neuron, 70(6), 1054–1069.

Sayers, K., & Menzel, C. R. (2012). Memory and foraging theory: Chimpanzee utilization of optimality heuristics in the rank-order recovery of hidden foods. Animal Behaviour, 84(4), 795–803.

Schuck, N. W., Cai, M. B., Wilson, R. C., & Niv, Y. (2016). Human orbitofrontal cortex represents a cognitive map of state space. Neuron, 91(6), 1402–1412.

Schultz, W., Dayan, P., & Montague, P. R. (1997). A neural substrate of prediction and reward. Science, 275(5306), 1593–1599.

Schwarz, G. (1978). Estimating the dimension of a model. The Annals of Statistics, 6(2), 461–464.

Seidel, D. P., & Boyce, M. S. (2015). Patch-use dynamics by a large herbivore. Movement Ecology, 3(1), 7.

Staddon, J. E., Hinson, J. M., & Kram, R. (1981). Optimal choice. Journal of the Experimental Analysis of Behavior, 35(3), 397–412.

Stephens, D. W., & Krebs, J. R. (1986). Foraging theory. Princeton: Princeton University Press.

Sutton, R. S., & Barto, A. G. (1998). Reinforcement learning: An introduction. Cambridge: MIT press.

Todd, I. A., & Kacelnik, A. (1993). Psychological mechanisms and the marginal value theorem: dynamics of scalar memory for travel time. Animal Behaviour, 46(4), 765–775.

Sweis, B. M., Thomas, M. J., & Redish, A. D. (2018). Mice learn to avoid regret. PLoS Biology, 16(6), e2005853.

Wilson, R. C., Takahashi, Y. K., Schoenbaum, G., & Niv, Y. (2014). Orbitofrontal cortex as a cognitive map of task space. Neuron, 81(2), 267–279.

Wittmann, M. K., Kolling, N., Akaishi, R., Chau, B. K., Brown, J. W., Nelissen, N., & Rushworth, M. F. (2016). Predictive decision making driven by multiple time-linked reward representations in the anterior cingulate cortex. Nature Communications, 7(1), 1–13.

Yoon, T., Geary, R. B., Ahmed, A. A., & Shadmehr, R. (2018). Control of movement vigor and decision making during foraging. Proceedings of the National Academy of Sciences, 115(44), E10476–E10485.

